# Down- to up-state transition is the default pathway in TREK K_2P_ channel activation and does not involve a lipid occluded pore

**DOI:** 10.64898/2026.05.07.723243

**Authors:** Marianne A. Musinszki, Chun Kei Lam, Edward Mendez Otalvaro, Friederike Schulz, Elena B. Riel, Anthony Ogwo, Kristin Rathje, Lea C. Neelsen, Bert L. de Groot, Marcus Schewe, Thomas Baukrowitz

**Affiliations:** Physiological Institute, Kiel University, Kiel, Germany; Department of Theoretical and Computational Biophysics, Max Planck Institute for Biophysical Chemistry, Göttingen, Germany; Department of Anesthesiology, Weill Cornell Medical College, New York

**Author notes:** shared last author (Marcus Schewe, Thomas Baukrowitz).

**Keywords:** TREK/TRAAK K_2P_ channels, selectivity filter gating, cysteine scanning mutagenesis, MD simulation, free energy calculation

## Abstract

Two crystallographic states of mechanosensitive TREK/TRAAK K_2P_ channels - a low-activity down-state and a high-activity up-state - have been proposed to underlie gating, but the origin of the low activity remains debated. Competing models suggest either lipid-mediated pore block or selectivity filter (SF) inactivation. Using systematic mutagenesis of M2/M4 helices, we identified 16 highly active mutants and assessed their activation mechanisms via free-energy calculations, molecular dynamics simulations, and a state-dependent pharmacological probe. The computational approaches reliably predicted mutation-induced shifts in the down-up equilibrium. We further show that intracellular acidification and regulatory lipids primarily stabilize the up-state, consistent with stretch, temperature, and dephosphorylation. These findings support the down-up transition as the principal physiological activation pathway and suggest that mechanosensitivity arises from the larger membrane footprint of the up-state. Our data argue against a physiological role of a lipid-blocked pore and instead support gating via conformational control of the SF in TREK/TRAAK channels.

## Introduction

Tandeml-pore domain potassium (K_2P_) channels generate background K⁺ currents that stabilize the resting membrane potential and dampen cellular excitability. TREK channels of the TREK/TRAAK subfamily are widely expressed in the central and peripheral nervous system and in peripheral tissues, making them important targets for pharmacological modulation. Their activity is strongly regulated by diverse physiological cues, including membrane stretch, temperature, intral- and extracellular pH (pH_i_ and pH_e_), membrane lipids, and (de-)phosphorylation^1–5^.

Crystallographic structures of TREK/TRAAK channels revealed two global conformations - termed “down” and “up” - that primarily differ in the orientation of the intracellular portions of helices M2/M4 and in the accessibility of intramembrane fenestrations between the two subunits that branch-off the central cavity. In the down conformation (down-state), these fenestrations provide lateral access from the membrane to the inner cavity, whereas in the up conformation (up-state), they are sealed by an upward movement and rotation of M4. The statel-dependent inhibitor norfluoxetine (NFx) binds within the fenestrations preferentially in the down-state and thereby can be used as conformational reporter^6–9^.

While mechanical activation is associated with a down-to-up transition, the mechanistic basis of the basal low-activity state remains controversial. One model, based mainly on structural data, proposes that in down-like conformations an acyl chain enters the cavity through the membrane-facing fenestrations and sterically occludes the permeation pathway^10–13^. However, other work emphasizes that K_2P_ channels gate predominantly at the selectivity filter (SF), which is highly conductive in the up-like conformation but inactivated in the down-like conformation. This view is supported by functional data and by molecular dynamics (MD) simulations showing that ion occupancy of the SF strongly controls channel activity independently of the conformational state of the lower helices^8,9,14–17^. In addition, the observation that the small-molecule activator 2-aminoethyldiphenyl borate (2-APB) can strongly activate TREK-1/-2 channels in the down-state is difficult to reconcile with a lipid-blocked pore^18^.

Various stimuli activate TREK/TRAAK channels and have been classified based on their NFx sensitivity. Mechanical stretch, dephosphorylation and temperature promote the up-state^7–10,15,16^, as indicated by reduced NFx sensitivity, whereas intracellular acidification and membrane depolarization activate channels that largely remain in the NFx-sensitive down-state^6,8^. For other activators, such as PIP_2_ or lysophospholipids, the underlying mechanism remains unclear.

Several gain-of-function (GOF) mutations have been identified in TRAAK and TREK-1/-2 and can be grouped according to whether they promote the up-state (up-state inducing mutants) or enhance activity while remaining in the down-state (down-state activators)^7,8,12,19,20^. Notably, the first published crystallographic structures of GOF mutants in TREK/TRAAK channels were initially interpreted to suggest that the down-state represents the physiologically active conformation^19^. However, because the up-state occupies a larger membrane footprint and, therefore, is promoted by membrane stretch, it is now generally considered the high-activity state, whereas the down-state represents the basal low-activity conformation in the unstretched membrane^7,11,12,16^.

In this study, we performed systematic scanning mutagenesis of the M2–M4 helices and identified 16 highly active mutants. Using a combination of free-energy calculations (FEC), all-atom MD simulations, and an NFx-based conformational assay, we determined whether mutations promote the up-state or enhance activity via alternative mechanisms. This combined approach of structural, electrophysiological and computational techniques provided mechanistic insight into the structural determinants governing the thermodynamic stability of the two crystallographic states and demonstrated that computational methods can robustly predict mutation-induced shifts in the down-up equilibrium. We further examined the mechanisms of further physiological activators (i.e., pH, PIP_2_ and regulatory lipids), and evaluated the physiological relevance of lipid occlusion in TREK/TRAAK channel gating.

## Results

### Scanning mutagenesis identified 16 highly active mutants in TREK-1

We conducted a systematic cysteine scanning mutagenesis comprising 81 positions along the transmembrane helices M2, M3 and M4 to identify critical residues and by studying these to better understand the gating mechanisms in TREK channels (Fig. 1a).

**Figure 1:**
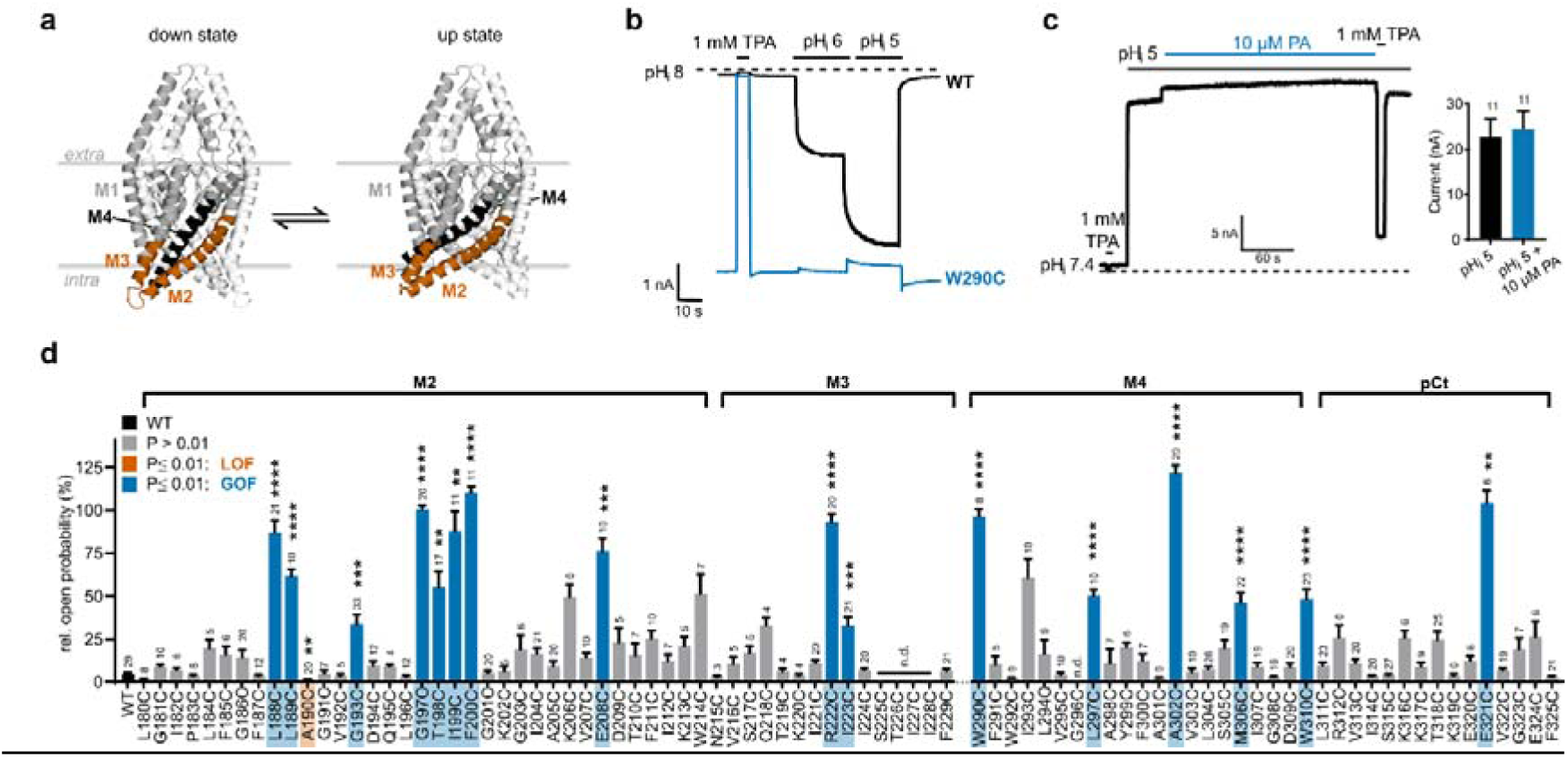
Systematic cysteine scanning mutagenesis of lower M2, M3 and M4 and determination of relative *P*_o_ in TREK-1 channels by pH_i_ activation. a TREK-2 crystal structure in the up- and down-state with the investigated regions of one subunit highlighted (M2/M3 orange, M4 black). **b** Representative measurements of wildtype (WT) TREK-1 channels and the W290C GOF mutant with high relative *P*_o_. Currents were normalized to the respective pH_i_ 5-activated current. **c** Representative current trace and bar graph showing the maximal activation of WT TREK-2 channels by intracellular acidification to pH_i_ 5. Additional application of 10 µM PA had little effect. **d** Relative *P*_o_ of TREK-1 WT and cysteine mutants derived from measurements as shown in c. Mutants with a relative *P*_o_ different from WT were classified as LOF or GOF phenotypes by Brown-Forsythe and Welsh one-way ANOVA with Dunnett’s T3 for multiple comparisons as post-hoc test (F(DFn, DFd) = 36.45 (82.00, 35.61), p < 0.0001; W(DFn, DFd) = 55.89 (82.00, 207.8), p < 0.0001, α = 0.01; adjusted P values). P-value symbols of statistical analysis are denoted next to the bars (**p ≤ 0.01, ***p ≤ 0.001, ****p ≤ 0.0001). All data shown were recorded in inside-out patches from *Xenopus* oocytes in symmetrical 120 mM K^+^ solutions at -80 mV with a continuous voltage protocol, and are presented as mean ± SEM with the number of experiments denoted next to the bars. n.d., not determined; pCt, proximal C-terminus.

We expressed TREK-1 channels in *Xenopus* oocytes and measured currents in inside-out patches. The basal activity of TREK-1 at high pH_i_ (e.g. 8) is very low but increases strongly (∼25-fold) upon acidification to pH_i_ 5 (maximal pH_i_ activation^2^) (Fig. 1b). Activation by intracellular acidification is so robust and strong that application of known additional activating stimuli (e.g. PIP_2_ or PA) produced only little further activation in TREK channels (Fig. 1c). Therefore, a mutation that raises (lowers) the basal activity is expected to reduce (increase) the margin for further channel pH_i_ activation and, thereby, can be classified as GOF (or loss-of-function, LOF) mutation. This type of assay has the advantage that it is insensitive to effects of the mutation on the expression level or intrinsic variations of expression related to the RNA quality or duration of expression.

With this rationale we used the extent of maximal pH activation (i.e. pH_i_ 5) relative to the current at pH_i_ 8 to calculate a relative open probability (relative *P*_o_). An example for a known GOF mutation (W290C) analysed this way showed very high basal currents and abolished pH_i_ 5 activation (Fig. 1b). pH_i_ 5 even produced a slight inhibition (probably conduction H^+^ block) resulting in a relative *P*_o_ apparently above 100 %).

We used variance analysis to determine which of the 81 mutants had a relative *P*_o_ different from WT TREK-1 channels that had a relative *P*_o_ of 4.8 ± 0.7 % (Fig. 1d). 64 mutations were not different from the WT (P ≥ 0.01) and only one mutation (A190C) led to a LOF phenotype with reduced relative *P*_o_ (1.6 ± 0.3 %). Notably, a total of 16 mutations were categorized as GOF mutations with relative *P*_o_ ranging from 33.0 ± 4.6 % to 121.9 ± 4.4 % (Table S1).

### FEC to predict the conformational state of TREK channels

If the crystallographic down-state and up-state structures of TREK-2 represent conformations that are similar to those that the channels actually adopt in the inside-out patches of our experiments, it might be possible to predict whether a mutation would cause a GOF/LOF phenotype *in silico*.

We approached this question by performing non-equilibrium FEC^21^ with the two crystallographic states of TREK-2 as starting structures^7^ since only a crystallographic up structure is currently available for TREK-1 channels. We used Amber14sb and CHARMM36m force fields to estimate the free energy differences ΔΔ*G* between the crystallographic up- and down-states of TREK-2 mutants relative to the WT (Fig. S1). Results from both force fields were then averaged to provide a consensus estimate ΔΔ*G*_c_ of the free energy difference^22^.

We performed this analysis for 12 of the GOF mutations and also included 4 mutations whose relative *P*_o_ were not different from the WT. Residues were grouped according to the ΔΔ*G*_c_ calculated for the corresponding TREK-2 residues (Fig. 2a, b and Table S2). We regarded a conformational shift towards the up-state or down-state conformation to occur when ΔΔG_c_ values were bigger or smaller than a RMS error of 4.2 kJ/mol^-1^, respectively (as benchmark for the accuracy of the FEC method^23–25^. For a group of 5 GOF mutants (G212C, F215C, A318C, M322C, and W326C) ΔΔ*G*_c_ were ≥ 4.2 kJ mol^-1^, suggesting a conformational shift towards the up-state conformation, consistent with the high relative *P*_o_. However, for the other 7 GOF mutants (L203C, L204C, G208C, T213C, I214C, W306C, and L313C) the ΔΔ*G*_c_ values predicted no change of the conformational equilibrium or even shifted it towards the down-state (≤ 4.2 kJ mol^-1^).

**Figure 2:**
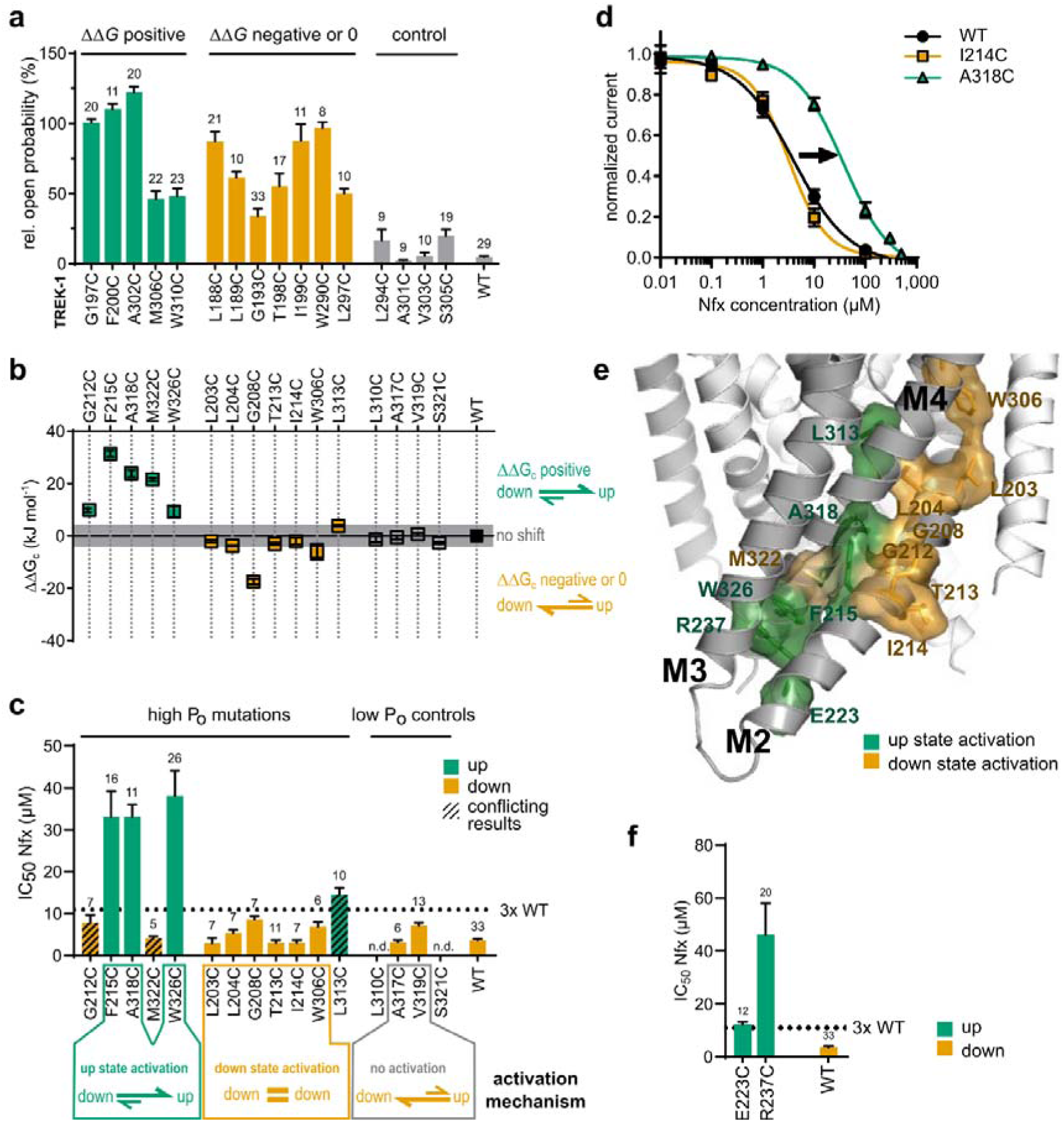
TREK up- and down-state equilibrium predicted by FEC and probed by state-dependent NFx inhibition. **a** GOF cysteine substitutions with high relative *P*_o_ and WT-like control residues grouped by the predicted shift of the conformational equilibrium according to ΔΔ*G*_c_ values from FECs shown in b. **b** ΔΔ*G*_c_ values from FECs of corresponding TREK-2 residues and WT-like control residues aligned to the upper panel (n = 10 for each mutation). ΔΔ*G*_c_ values ≥ 4.2 kJ mol^-1^ indicate destabilization of the down-state. **c** IC_50_ values of NFx inhibition for GOF cysteine substitutions with high relative *P*_o_ but different ΔΔ*G*_c_ values. Mutants adopting the up-state are coloured green, mutants in the down-state are coloured orange. The labelling below describes the determined activation mechanism for the groups. **d** Exemplary NFx dose-response relationships for a mutant in the down-state (I214C) and a mutant in the up-state (A318C) compared to WT TREK-2 channels. Data were recorded in inside-out patches of *Xenopus* oocytes in physiological K^+^ gradients with a ramp protocol, and normalised mean currents + SEM at +40 mV were plotted (n = 7-33 for each concentration). **e** Positions of highly activating mutations upon cysteine substitution in M2 and M4 shown in the down-state structure of TREK-2. **f** IC_50_ values of NFx inhibition for cysteine mutants with high relative *P*_o_ at charged residue positions not amenable to FECs. All data are presented as mean ± SEM with the number of experiments denoted next to the bars.

Thus, for these mutations the FEC either failed to generate a meaningful prediction or the mutation indeed did not substantially change the down-up equilibrium but caused activation by an alternative mechanism (i.e. not involving an up-state transition). To discriminate between these alternatives, we utilized NFx, a state-dependent inhibitor that binds preferentially to the down-state^7^.

### Using a pharmacological probe to evaluate the GOF contribution of the up-state conformation

For the basal activity of WT TREK-1 channels we measured the NFx sensitivity (IC_50_ of 3.7 ± 0.3 µM) indicating that the channels adopted mostly the low activity down-state conformation. We then determined the NFx sensitivities of the 12 GOF mutants investigated by FECs and also included two control mutants (A317C and V319C) with WT-like relative *P*_o_ and ΔΔ*G*_c_ estimates (Fig. 2c and Table S2). We categorized the mutants as WT-like if the apparent affinity for NFx was not more than 3-fold different from the WT. Accordingly, larger shifts to higher values indicated that the channels to a substantial degree adopted an up-state like, NFx insensitive conformation.

In the group of five mutations with high relative *P*_o_ and a predicted conformational shift to the up-state according to the FECs, the NFx affinities of three mutations (F215C, A318C, W326C) were strongly (i.e. about 10-fold) reduced compared to the WT (IC_50_ 33.1 ± 6.2 µM, 33.1 ± 2.2 µM, and 38.1 ± 6.1 µM, respectively), consistent with the predicted conformational shift towards the up-state conformation. In contrast, G212C and M322C apparently did not substantially adopt the up-state conformation, as the NFx sensitivities were WT-like (IC_50_ 7.8 ± 1.9 µM and 4.1 ± 0.5 µM) and, thus, these results were inconsistent with the FEC predictions. We must therefore assume that the mutations activated the channel without inducing the up-state (down-state activating mutation).

In the group of seven high relative *P*_o_ mutants predicted to adopt mostly a down-state conformation, six mutants showed indeed little change in NFx sensitivity as predicted by the FEC suggesting that their conformation resembles the down-state conformation. Only L313C exhibited an intermediate NFx sensitivity (IC_50_ 14.6 ± 1.5 µM) indicating some contribution of the up-state. However, the ΔΔ*G*_c_ value (3.78 ± 0.39 kJ mol^-1^) was just below to the detection threshold consistent with some shift to the up-state conformation. Furthermore, the two control mutations (A317C and V319C) did not change the NFx sensitivity consistent with FEC prediction. Therefore, for 11 mutations (six down-state activation mutants, three up-state inducing mutants and two control mutants) the FEC correctly predicted whether a mutation would promote the up-state conformation. Only for three mutants the FEC predictions were ambiguous as they were not consistent with observed NFx sensitivities. Thus, overall, the FEC predictions are in good qualitative agreement with the pharmacologically assessed direction of a shift in the down-to-up state equilibrium.

### Investigation of the up/down equilibrium with unrestrained MD simulations

We next examined whether unrestrained MD simulations can also predict the effects of point mutations on channel conformation. We performed multiple 1 µs all-atom simulations of WT channels, four up-state inducing mutations (F215C, R237C, A318C, W326C), the three in FEC and NFx conflicting mutants (G212C, L313C, M322C) as well as for two WT-like controls (A317C, V319C) (Fig. 2 c, f). However, not all up-state inducing mutations transitioned to the up-state during all runs within the limited simulation time (Fig. S2). Thus, we analysed the conformational sampling using principal component analysis and extracted Eigenvectors (EV) that capture the high-variance motions during conformational change. EV1 and EV3, that describe movements of the M2-M4 helices, were used to visualize their conformational spaces in the up- and the down-state as contour plots (Fig. 3a).

**Figure 3:**
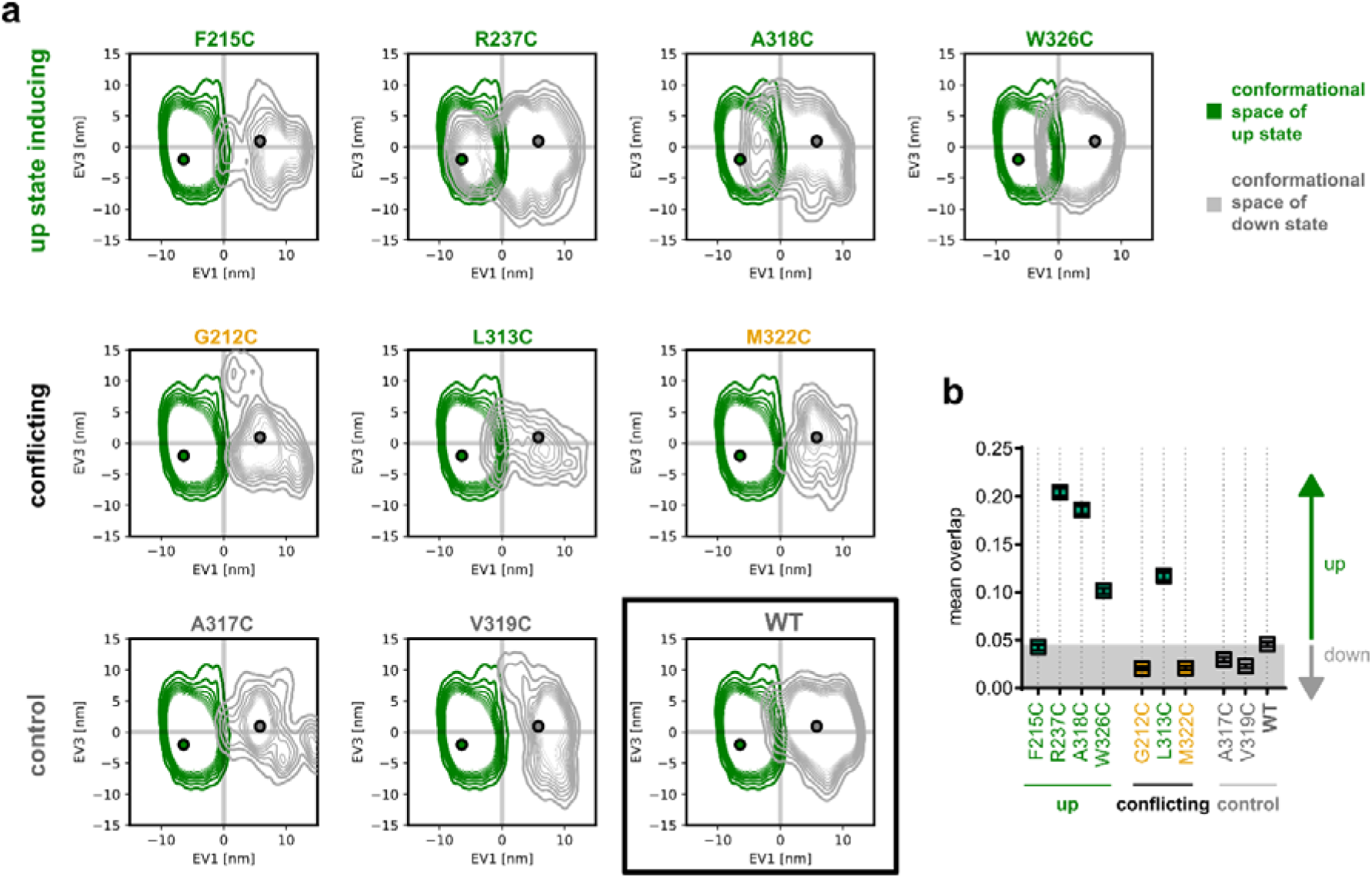
Analysis of conventional MD simulations to characterise up-state inducing mutations. **a** Contour plots of EV1 and EV3 which represent the conformation of lower M2/M3/M4 helices, showing the conformational spaces of the WT and different mutants in the down-state (grey) and the up state (green). **b** Mean overlap of the conformational spaces of down and up state in TREK WT and mutant channels derived from the contour plots in a. Green text marks mutants classified as up-state activating in the functional Nfx assay, orange text marks down-state activating mutants (G212C and M322C), and grey marks WT and controls (A317C and V319C).

We quantified the mean overlap between down- and up-state conformational spaces as a measure of the probability of state transitions (Fig. 3b, Table S3). In WT and WT-like controls (A317C, V319C), conformational spaces remained largely separated. In contrast, three up-state inducing mutants (R237C, A318C, W326C) showed a pronounced shift of the down-state ensemble toward the up-state space, increasing overlap and indicating destabilization of the down-state. Notably, F215C did not show increased overlap despite its functional phenotype. We further analysed mutants with discordant FEC and NFx classifications (G212C, L313C, and M322C; Fig. 2c). In all cases, the MD-based analysis was in agreement with the functional NFx results, showing increased overlap for L313C (up-state) and reduced overlap for G212C and M322C (down-state).

To gain insight into the structural basis for up-state transition, we additionally analyzed the residue interaction networks for the up-state inducing mutants and WT-like controls (Tables S4, S5). The contact network of WT-like A317C and V319C was relatively unchanged. F215, despite the down-like overlap, displayed a tendency towards the up-state. F215 has an extended contact network in the WT that is known to stabilize the M2-M4 bundle^7,16^ (21 interactions in total, i.e. in both subunits), and we observed a collapse of hydrophobic contacts with a loss of all interactions (π-π stacking and van der Waals) to F244 in M2 and Y315 in M4 upon cysteine substitution. Furthermore, as these residues encompass the fenestration, structural changes would be consistent with a high relative *P*_o_ and reduced NFx sensitivity. W326 and R237 side chains are stacked with M322 in the WT down-state, which also stabilizes the lower M2/M3 bundle^7,16^. Upon mutation, mutual cation-π interactions are lost, which enables M2/M4 dissociation. The bulkier cysteine in A318C led to steric clashes with the opposed M2 and close contacts to F215, L211 and G208 of the GXG motif that permits M2 kinking and is thought to be coupled to the rearrangement of M4 during up state transition^19^, making it conceivable that local deformation of M2 induces evasion of M4 towards the up-state conformation. L313C showed only slightly less van-der-Waals contacts, but, in contrast to the other up-state inducing mutants, rather large protein-wide shifts in intra-chain H-bonds, inter- and intrachain π-stacking, ionic and van-der-Waals interactions as well as the loss of all π-cation interactions. Therefore, the L313C substitution may differ in the mechanism that shifts the conformational equilibrium to the up-like state.

Overall, these data indicate that the down-state is more sensitive to mutational perturbation and demonstrate that MD-based conformational analysis reliably predicts mutation-induced shifts in state equilibrium, even in cases where FEC results are ambiguous.

### Systematic substitutions at A318 reveal steric control of the down/up equilibrium

To further extend our approach, we systematically examined substitutions at position A318. In our cysteine scan, A318C displayed the highest relative *P*_o_ and strongly destabilized the down-state in FEC (ΔΔ*G* = 23.84 ± 1.29 kJ mol⁻¹). We therefore tested whether the effects of substitutions differing in side chain volume could be predicted (G < C < V ≈ N < F < Y).

Consistent with this expectation, down-state destabilization predicted by FEC increased with side chain volume, ranging from A318G (ΔΔ*G*_c_ = 7.51 ± 0.22 kJ mol⁻¹) to A318Y (ΔΔ*G*_c_ = 101.89 ± 3.02 kJ mol⁻¹; Fig. 4a, Table S6). Electrophysiology confirmed the high relative *P*_o_ for bulkier substitutions, whereas A318G produced a LOF phenotype with reduced basal current and relative *P*_o_ Fig. 4a, Table S7. NFx assays further supported these assignments: A318F exhibited a >50-fold increase in IC₅₀ (155.7 ± 7.0 µM), indicative of up-state stabilization, whereas A318G showed markedly increased NFx sensitivity (IC₅₀ = 0.2 ± 0.1 µM), consistent with a down-like conformation (Fig. 4b, c).

**Figure 4:**
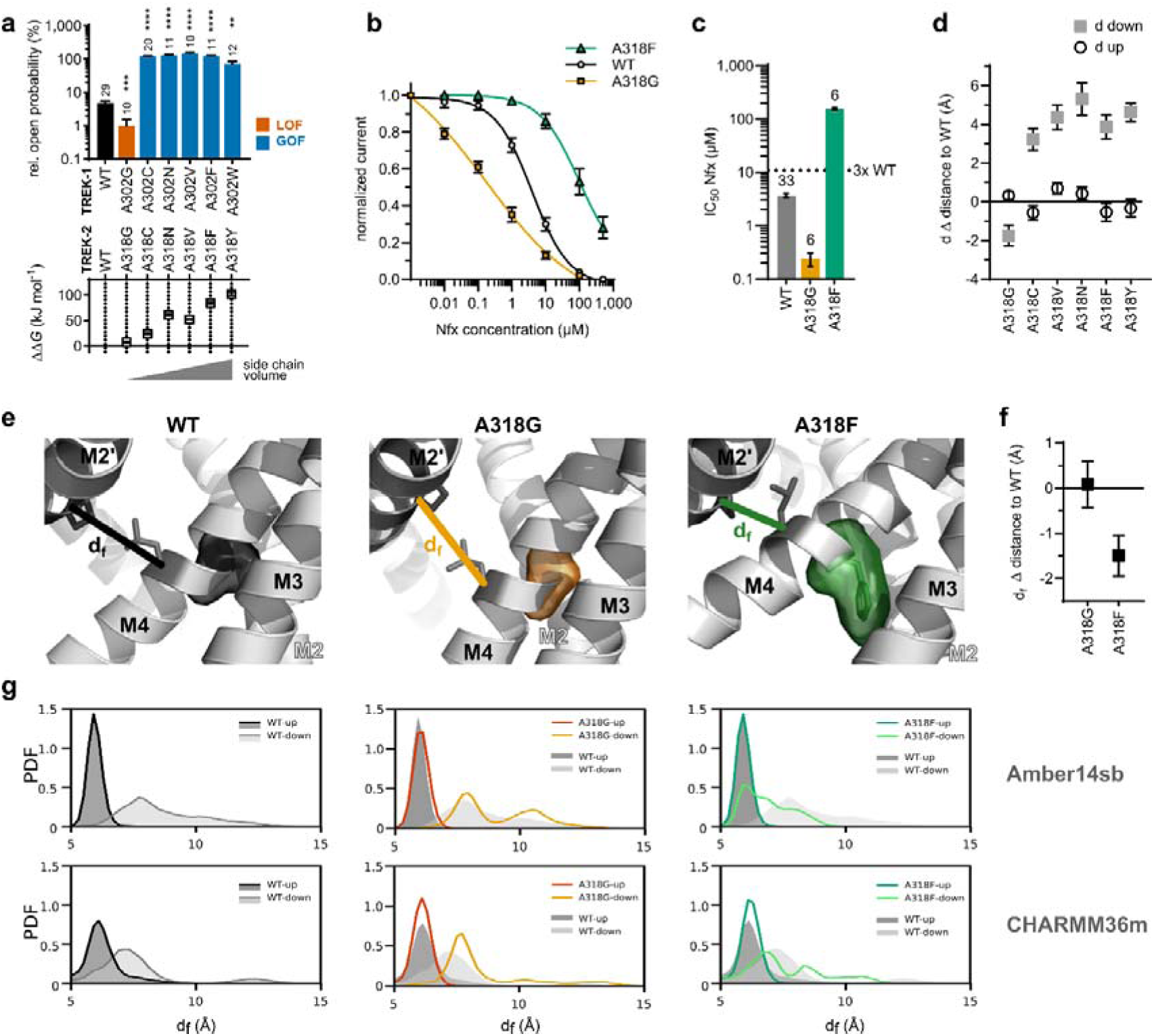
Different substitutions at residue A318 probed by electrophysiology and computational methods reveal an up-state activation mechanism and a contracted down-state in TREK channels. **a** Relative *P*_o_ of different substitutions at A302 in TREK-1 channels and ΔΔ*G* values from FECs for the corresponding TREK-2 A318 residue (Brown-Forsythe and Welsh one-way ANOVA with Dunnett’s T3 post-hoc test (F = 116.8, W = 311.5, α = 0.05; **p ≤ 0.01, ***p ≤ 0.001, ****p ≤ 0.0001). Note that for technical reasons Y was simulated instead of W. **b** NFx dose-response-relationships for A318G and A318F substitutions measured in inside-out patches of *Xenopus* oocytes in physiological K^+^ gradients with a ramp protocol, and normalized mean currents at +40 mV were plotted. **c** IC_50_ concentrations derived from measurements in b. Data are presented as mean ± SEM with the number of experiments denoted next to the bars. **d** Mean distance difference dΔ to the WT in the down- and up-state for different A318 substitutions. **e** Fenestration site of TREK-2 WT, A318G, and A318F channels, showing the location of the substitution and the d_f_ distance measurement used to assess fenestration diameter. **f** Distances d_f_ in the A318G and A318F mutants derived from MD simulations. **g** Probability density functions (PDF) for d_f_ distances of A318G and A318F compared to the WT (shown as shadow) with both force fields used, taken from the last 500 ns of simulations. Only means are shown for clarity, data presented with 95 % confidence intervals can be found in Fig. S3.

MD simulations revealed that larger side chains introduce steric clashes that disrupt tight packing of the M2–M4 helix bundle, promoting transition to the up-state as shown by the shifted probability density functions (Fig. 4g). Accordingly, the M2-M4 distance measured between C_α_ of G216 (M2) and W326 (M4) was increased in all up-state inducing mutants, but reduced in A318G, which stabilized a more compact down-like conformation (Fig. 4d, g Table S8). Analysis of the fenestration geometry (as the distance d_f_ between C_α_ of P198 and L320) showed that A318G widens the fenestration despite overall compaction of the helix bundle, likely enhancing NFx accessibility and/or binding (Fig. 4e, f). Together, these data identify A318 as a key determinant of the down/up equilibrium, where side chain volume controls conformational transitions through modulation of helix packing.

### Physiological stimuli exclusively promote the up-state

We identified at least five mutations that promote transition to the up-state, demonstrating that this conformation is intrinsically stable and can be adopted independently of membrane stretch. Consistently, dephosphorylation and temperature have previously been shown to favour the up-state^8,9^. We therefore examined whether additional physiological stimuli, i.e. intracellular acidification (pH□), PIP_2_, (lyso-)phospholipids (LPA, PA), and extracellular LPC, induce similar transitions (Fig. 5a).

**Figure 5:**
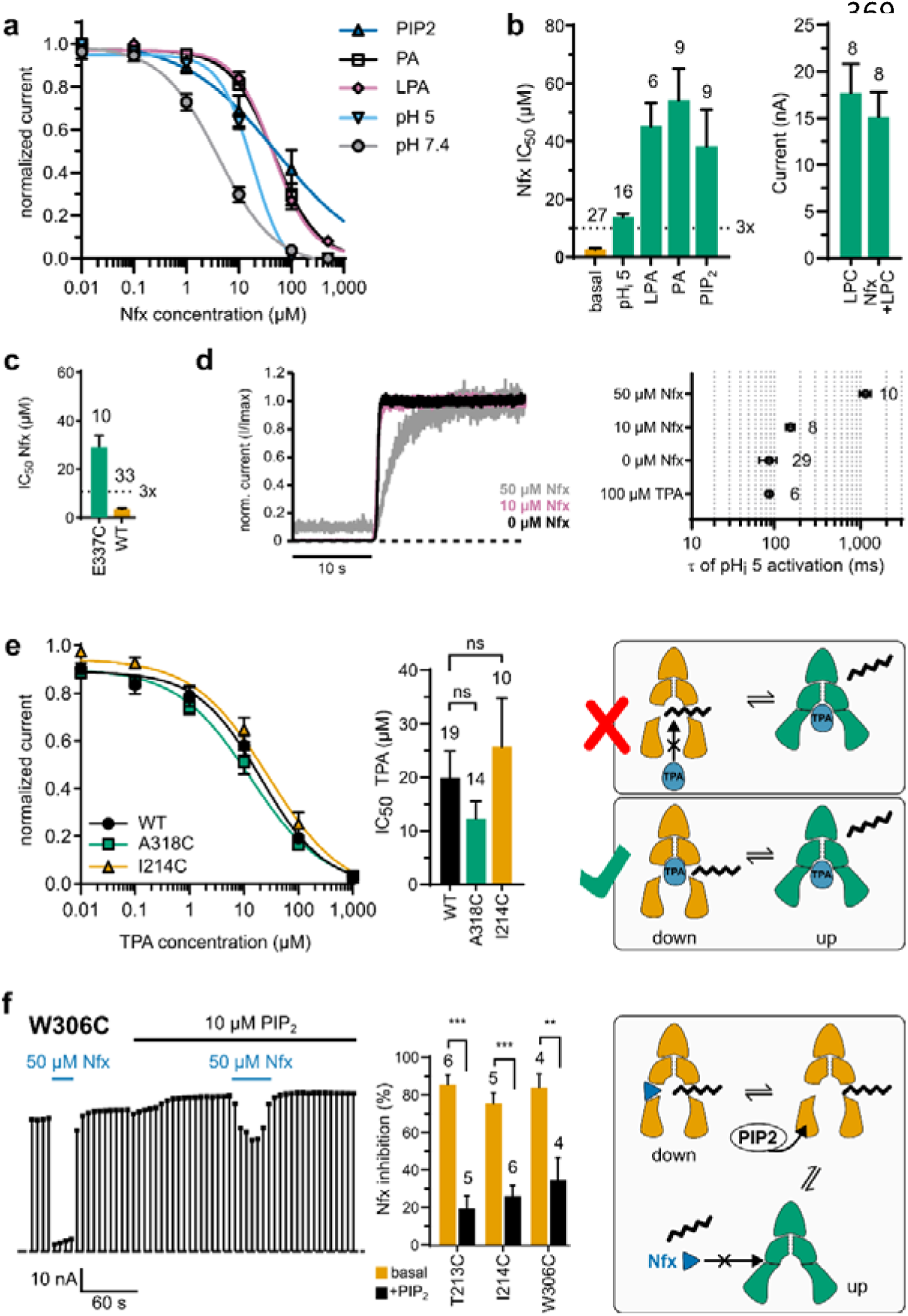
Up-state activation by physiological stimuli and absence of a ’gating lipid’. **a** NFx dose-response relationships for WT TREK-2 in basal conditions and in the presence of different physiological activators. Data were recorded in inside-out patches of *Xenopus* oocytes in physiological K^+^ gradients with a ramp protocol, and normalized mean currents + SEM at +40 mV were plotted (n = 3-21 for each concentration). **b** Left, IC_50_ values of NFx inhibition derived from the dose-response relationships in a. Right, activation of TREK-1 channels by extracellular application of 10 µM LPC in the absence and presence of 10 µM NFx recorded in transiently transfected HEK293 cells. **c** IC_50_ values of NFx inhibition for E337C compared to the WT. **d** Left, exemplary recordings of the time course of pH_i_ 5 activation of TREK-2 in the presence of 100 µM TPenA or different NFxconcentrations (0 - 50 µM). Currents were normalized to the maximal current for each condition. Right, τ of activation by pH_i_ 5 derived from mono-exponential fits for each condition. **e** Left, TPenA dose-response relationships for WT TREK-2, the up-state inducing mutant A318C, and the down-state activating mutant I214C. Data were recorded as in a. Right, cartoon illustrating the lack of competition between TPenA and an acyl chain in both mutants. **f** Exemplary recording of the NFx inhibition of the down-state activating TREK-2 mutant W306C before and during application of 10 µM PIP_2_. Data were recorded as in a, and the current at +40 mV was plotted over time. Middle, percent inhibition by NFx before and during PIP_2_ activation for three down-state activating mutants (paired two-tailed *t*-test with Holm-Sidak correction for multiple comparison). Right, cartoon illustrating the reduced NFx inhibition when the down-state activating mutant is converted to the up-state by PIP_2_.

Intracellular acidification increased the NFx IC₅₀ ∼5-fold (to 13.9 ± 1.1 µM), whereas LPA, PA, and PIP_2_ produced larger shifts (14 - 20-fold; IC₅₀ = 45.2 ± 8.1 µM, 54.2 ± 10.8 µM, and 38.4 ± 12.6 µM, respectively; Fig. 5b; Table S10). Extracellular LPC similarly rendered channels largely insensitive to NFx (Fig. 5b; Fig. S4). These findings indicate that diverse physiological stimuli robustly promote the up-state.

The comparatively modest shift observed for acidification suggests that a fraction of channels remains in the down-state at pH□ 5. As activation is mediated by protonation of E337, incomplete protonation likely limits full transition. Consistently, neutralization of E337 (E337C) further reduced NFx sensitivity (IC₅₀ = 30.2 ± 3.7 µM; Fig. 5c), indicating enhanced stabilization of the up-state. To further evaluate whether pH□-induced activation involves an up-state transition, we analysed the kinetics of rapid pH□ 5 activation in the presence of NFx (0 - 50 µM; Fig. 5d). Indeed, NFx markedly slowed channel activation, increasing the time constant (τ) by ∼10-fold at 50 µM, consistent with state-dependent binding and indicating that NFx dissociation precedes the transition to the up-state. In contrast, the pore blocker TPenA, which binds to basal and activated states of TREK-1 with similar affinity^26^, did not affect activation kinetics, demonstrating that the slowing is specific to NFx.

Taken together, these findings demonstrate that physiological stimuli promote TREK activation predominantly by stabilizing the up-state.

### High-activity down-state mutations argue against a lipid-block mechanism

Two models have been proposed to account for the low activity of the down-state in TREK/TRAAK channels: pore occlusion by lipids^11,12^ vs inactivation at the SF^8,9,14,15,18^. We identified multiple GOF mutants that display high activity while retaining strong NFx sensitivity, indicating a predominant down-state conformation (Fig. 2).

A lipid occupying the pore in the down-state would be expected to compete with cavity blockers such as TPenA, leading to a lower apparent affinity compared to a lipid-free up-state. However, the apparent TPenA affinity was not different in an up-state inducing mutant (A318C), in the WT channel in the basal state, or in a down-state activating mutant (I214C; Fig. 5e, Table S11).

Furthermore, if lipid occlusion were a major contributor to the low activity in the down-state, stabilization of the up-state - for example by PIP_2_ - should relieve lipid occlusion and produce an additional current increase also in the down-state activating mutants. We used the W306C mutant (Fig. 1b), as substitutions at this position are thought to reside in the down-state and activate the SF directly^8,19,20^, as also shown in this study. Application of PIP_2_ produced only a very modest current increase (17 ± 10 %), while strongly shifting the conformational equilibrium toward the up-state, as evidenced by a marked reduction in NFx inhibition (from ∼84 % to ∼34 %; Fig. 5f, Table S12).

## Discussion

In this study, we identified gating relevant positions which, upon mutation, strongly activate TREK channels. Using a combination of computational and functional approaches we predicted their conformation, evaluated *in silico* structure prediction approaches and gained insight into the activation mechanisms of TREK channels.

### Low-cost free energy simulations robustly predict mutation-induced conformational states

We initially took advantage of FEC derived from non-equilibrium MD simulations as a fast method to predict the conformational shifts brought by mutations using structural data for distinct functional states, a method to date used mainly for ligand-binding studies^22,27,28^. Our results show that this method had a robust predictive power that for ∼79 % of mutants assigned the correct conformational state and allowed the discrimination of the underlying activation mechanism in 11 of 14 high relative *P*_o_ mutants. It separated mutations inducing up-state transition from alternative down-state activation mechanisms that were revealed by the functional NFx assay. It further correctly predicted the effect of six point mutations at position A318, and therefore is a suitable tool with low computing costs for the initial investigation of distinctive point mutations in ion channels. Conventional MD simulations added detailed insight into structural rearrangements caused by distinct mutations with a focus on the lower M2/M3/M4 helix bundle, and enabled the investigation of residues either more challenging to converge in FEC (such as charge changes) or conflicting with functional measurements. MD observations using analysis of gating relevant Eigenvectors matched the results for up-state inducing and down-state activating mutants that had been assigned by FEC and functionally confirmed by the NFx assay. In the cases of three conflicting results (G212C, M322C, and L313C), MD simulations were consistent with the functional measurements, outperforming the faster FEC method.

### Mutation scanning reveals different networks for down- and upstate activation

Interestingly, our comprehensive scanning mutagenesis of TREK-1 channels revealed only one LOF mutant, while a comparably high number of mutations (16 out of 86) led to an activated phenotype. About half of the highly active mutants amenable to FEC appeared to preferentially adopt the down conformation similar to WT channels in the basal state. Down-state activation has been reported previously for several GOF mutants, i.e. G167I, G201D, and W306S (respectively G124I, G158D and W262S in TRAAK channels)^8,12,19^. Our study identified five more down-state activating mutations (plus W306*C*), highlighting that a down-state activating mechanism upon substitution is more prevalent than previously acknowledged. Functional and computational studies have shown that mutations close to the SF (e.g. W306C) can directly modulate SF gating, probably without a need for conformational change in the lower helices^18,20,29^. Our results, however, emphasize that down-state activation can be induced by mutations in the lower helix segments very distant from the SF. Down-state activating positions (L203, L204, G208, G212, T213, I214) formed an extended cluster in middle M2, with the exception of M322 on M4.

These residues were well interconnected in both up- and down-state, suggesting that a mutation here might destabilize both states about equally and, therefore, the channel remains in the down-state. How these mutations produce activation remains enigmatic. We speculate that the loss of interactions with M4, which in the WT channel occurs upon M4 upwards movement towards the up-state, are critical to induce activation of the SF. Correspondingly, the G208 position was described as a hinge that allows M2 buckling during up-state transition in TRAAK^19^, and G212 was recently identified as an effector site for activation by volatile anaesthetics, which disrupted M2/M3 interactions by steric crowding^30^.

Up-state inducing positions were located in the M2/M3/M4 bundle, and analysis of structural variances and interaction networks showed up-like conformations and the loss of contacts between M2 and M4 helices. For F215C, R237C and W326C, loss of stacked hydrophobic interactions led to up-state transition, in agreement with the crystallographic structure and subsequent MD studies^7,16^.

We find that this interaction core is more extensive and also comprises E223 (contacting R237) and A318. A318 represents a highly gating sensitive residue, because increasing M2/M4 distance by mutation predictably induced up-state transition, while tighter M2/M4 packing was seen in the A318G LOF phenotype, showing an even more compact ‘down+’-state in MD simulations as well as the NFx assay. While we cannot determine if the higher NFx affinity of A318G results from better access to the widened fenestration, improved binding, or from more channels being in the down- or ‘down+’-state, MD and functional results are consistent with the existence of a ‘super-down’-state as described by the computational study of Zhang *et al.* ^20^, and implies that a continuum of states around the ‘crystallographic snapshots’ exists.

### Up-state stabilization is the default physiological activation pathway

In TRAAK, increased membrane stretch induces the down-up transition, a mechanism also observed for TREK-2 in MD simulations□□¹□ and mimicked by arachidonic acid (AA)□. Increased temperature likewise promotes the NFx-insensitive up-state^8^. Intracellular acidification, reproduced by substitution of the pH sensor (E306 or E321, respectively, in TREK-1a and Trek-1b isoforms), similarly induces an up-state transition, as indicated by reduced fluoxetine sensitivity²□. In addition, we recently showed that (de)phosphorylation uses up-down transitions as a gating pathway to relay C-terminal signals to the SF^9^.

These observations raise the question of whether physiological stimuli generally activate TREK/TRAAK channels via a common mechanism. Our data indicate that this is indeed the case. All physiological activators tested - including intracellular acidification (or neutralization of the pH sensor), PIP_2_, PA, LPA, and extracellular LPC - shifted the conformational equilibrium toward the up-state. While McClenaghan *et al.*¹² reported down-state occupancy at pH□ 6, we observed a clear increase in NFx IC_50_ and a slowing of activation kinetics at pH□ 5 in the presence of NFx, consistent with up-state stabilization. This discrepancy is likely explained by incomplete activation at pH□ 6 and the limited sensitivity of the NFx assay when only a minority of channels occupy the up-state.

Together, these findings demonstrate that diverse physiological stimuli converge on a common gating mechanism by stabilizing the up conformation, establishing the up-state as the dominant functional state underlying TREK/TRAAK channel activation. A notable exception appears to be voltage-dependent activation, which can directly increase SF conductance without requiring an up-state transition^14,18^.

### Is mechanosensitivity in TREK/TRAAK channels physiological or epiphenomenal?

The finding that up-state transition represents the default pathway of physiological activation in TREK/TRAAK channels might reframe the role of mechanosensitivity. Rather than constituting a primary physiological gating mechanism, mechanosensitivity may reflect an experimental epiphenomenon arising from the larger membrane footprint of the up-state, which is stabilized by diverse physiological stimuli independent of membrane stretch. This interpretation provides an explanation for several puzzling observations. Although TREK/TRAAK channels display pronounced stretch sensitivity in excised membrane patches and reconstituted bilayers, mechano-activation is markedly attenuated in intact cells. Moreover, genetic ablation has not been associated with consistent reductions in mechanosensitivity at the cellular or tissue level. This contrasts with bona fide mechanosensitive channels such as PIEZO1/2, where a LOF results in well-defined mechanosensory deficits^31–33^.

### Pore occlusion by a gating lipid is unlikely to play a physiological role

It has been proposed that in TREK/TRAAK channels, the down-state is characterized by lipid occlusion of the pore, as electron density representing an alleged acyl was present at the fenestration in the down-state but not in the up-state^11,12^. However, this model is not supported by several lines of functional data reported previously as well as in this study.

First, it was shown that Rb⁺ ions strongly activate TREK/TRAAK channels when driven into the SF by membrane depolarization^8,14,34^. Importantly, this robust effect is restricted to the down-state, whereas membrane stretch or AA - both of which promote the up-state^7^ - strongly reduce Rb⁺-induced activation (Fig. S5). These findings imply that (i) Rb⁺ has unobstructed access to the SF in the down-state, inconsistent with a lipid-occluded pore, (ii) the SF is in a low-conductance (inactivated) configuration in the down-state, explaining the strong voltage-dependent activation by Rb⁺, and (iii) the SF is in a high-conductance configuration in the up-state, explained the lack of additional activation by Rb⁺.

Second, mutations within the SF, such as T212C at the S4 K⁺ binding site in TRAAK, alter ion occupancy and abolish activation by AA or mechanical stress^14^. This strongly supports the SF as the primary gating element. These findings are difficult to reconcile with a lipid-occlusion model, as it is unclear how marginal structural perturbations at the S4 binding site would affect the presence of the lipid that depends on a global conformational transition between the down- and up-state.

Third, we here report several GOF mutations located distal to the fenestrations that strongly activate TREK channels without promoting the up-state. These mutations are unlikely to alter lipid occupancy since they are not located in or at the fenestrations. Further, the pore blocker TPenA - expected to compete with a lipid for cavity binding - displayed similar apparent affinity in channels with open vs closed fenestrations.

Finally, we tested here directly the relative contributions of a putative lipid-occlusion and the SF gating using the W306C mutant, which is thought to act directly on the SF. W306C channels exhibit a high relative *P*_o_ and retain strong NFx sensitivity, indicating open fenestrations. If a lipid occludes the pore by binding to the fenestration in the down-state, closure of the fenestrations (e.g., in response to the up-state activator PIP_2_) should relieve this occlusion and produce a substantial increase in current similar to that observed in WT channels. However, PIP_2_ induced only a minimal increase in current, while clearly reducing NFx sensitivity, suggesting that fenestrations closed without removing a substantial pore occlusion. These findings do not support a major physiological role for lipid-occlusion in TREK/TRAAK gating. Instead, they are consistent with a model in which gating is primarily governed by conformational changes that regulate ion conduction at the SF (Fig. 6).

**Figure 6:**
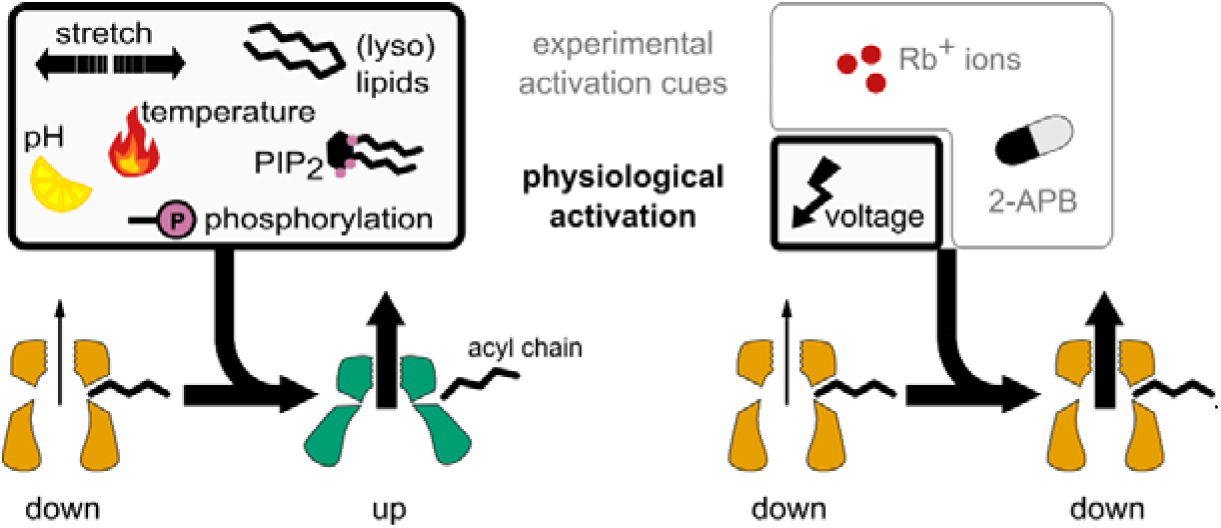
Proposed physiological gating mechanism. The cartoon on the left depicts how cellular stimuli including stretch and temperature induce the transition from the down-state with a lipid-free pore to the high activity up-state. The left cartoon shows SF activation of the lipid-free down-state by voltage or experimental cues. Physiological stimuli are highlighted bold.

## Methods

### Computational Methods

#### System construction

The up (PDB ID: 4BW5) and the down (PDB ID: 4XDJ) crystal structures of TREK-2 were used as the starting structures for the MD simulations^7^. Missing loops and residues were introduced using LOOPY^35^ and PyMOL^36^. The resulting structures were truncated at Lys333. CHARMM-GUI^37–39^ was used to embed TREK-2 into a 1-palmitoyl-2-oleoyl-sn-glycero-3-phosphocholine (POPC) membrane bilayer. The systems were solvated and neutralized by a 500 mM KCl solution. Two force fields, Amber14sb^40^ and CHARMM36m^41^ were used. For Amber14sb, Slipids^42^, the TIP3P water model^43^, and Joung and Cheatham ion parameters^44^ were used. For CHARMM36m, CHARMM36 lipids^45^, CHARMM36 TIP3P water model^46^ and CHARMM36 ion parameters^47^ were used. No membrane voltage was applied. An integration timestep of 2 fs was used for all simulations. All simulations were performed in GROMACS 2019 or GROMACS 2020^48,49^.

#### Free energy calculation (FEC)

Mutants that carried hybrid residues for the FEC were generated with pmx^50^. 10000 energy minimization steps were performed with a steepest descent algorithm preceding the simulations. Harmonic position restraints with a force constant of 1000 kJ mol^−1^ nm^−2^ were applied to all backbone atoms of the channel. All bonds were constrained with the LINCS algorithm^51^. As position restraints were applied to backbone C_a_ atoms of TREK-2, the free energy estimates represent the conformational shifts of restrained mutants.

For CHARMM36m simulations, Newton’s equations of motion were integrated with a leap-frog stochastic dynamics integrator at 300 K. Pressure was coupled with the Parrinello–Rahman barostat^52^ at 1 bar. Van der Waals forces were switched smoothly to zero between 1.0 to 1.2 nm. The particle mesh Ewald (PME) method^53^ with a 1.2 nm distance cutoff was used for electrostatic interactions. For Amber14sb simulations, a leap-frog algorithm was used as the integrator. Temperature and pressure were kept at 300 K with the velocity rescaling algorithm^54^ and 1 bar with the Parrinello–Rahman barostat. A cutoff of 1.5 nm was used for van der Waals interactions. The PME algorithm with a 1.5 nm distance cutoff was used for electrostatic interactions.

For every combination of systems, ten 20-ns equilibrium simulations were carried out. For the non-equilibrium simulations, starting snapshots were taken from the equilibrium runs every 80 ps (the first 1 ns was discarded). During the non-equilibrium simulations as TREK-2 is a dimer, two residues were morphed into target ones alchemically in 100 ps. The work values associated with the non-equilibrium transitions were computed via thermodynamic integration (TI). Combining the work values computed from the forward and the backward transitions via the Crooks fluctuation theorem^55^ and the Bennett Acceptance Ratio (BAR) method^56^ yielded the free energy differences ΔG_up_^mutation^ and ΔG_down_^mutation^. Point estimates of the free energy differences were obtained by including all the computed forward and backward work values. Uncertainties were estimated as the standard error of the ten independent estimates of ΔG, each coming from a unique pair of the distributions of forward and backward work values. For CHARMM36m calculations, ΔG_up_^mutation^ and ΔG_down_^mutation^ for glycine mutants were not provided because it is not yet possible to take the contributions of the grid-based energy correction maps (CMAP) into account when computing 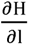 in GROMACS, and that the CMAP for glycine is different from those of other amino acids. As a result, the CHARMM36m free energy estimates for glycine mutants calculated with the current protocol would be invalid.

#### Conventional MD Simulations

Mutants for 1-μs unbiased simulations were generated with PyMOL^36^. The simulation setting was identical to that of the FEC except for the following: only bonds associated with hydrogen atoms were constrained with the LINCS algorithm in the conventional MD simulations. For CHARMM36m simulations, a leapfrog algorithm was used as the integrator, and the Nosé-Hoover thermostat^57,58^ and the Parrinello-Rahman barostat^52^ were used to maintain the systems at 300 K and 1 bar, respectively. For every combination of systems, 10000 energy minimization steps with the steepest descent algorithm followed by 4 to 10 independent 1-μs equilibrium simulations were carried out.

#### Difference Vector Projections

The difference vector between the up and down states was constructed from the two crystallographic conformations using the GROMACS module COVAR. Only C_α_ atoms of TREK-2 were used for the construction. The conventional MD simulation trajectories were then projected onto the difference vector using the GROMACS module ANAEIG. The projection values were re-centered and re-scaled such that the crystallographic up and down conformations have a difference vector projection value of 1 and -1, respectively.

#### Distance Profiles

For distance measurements between M2 and M4, the distance between C_α_ atoms of G216 and W326 were measured. For the fenestration, d_f_ was defined as the intersubunit distance between C_α_ atoms of P198 and L320. The distance means were averaged over all Amber14sb and CHARMM36m simulations. Errors are calculated as the standard deviation of the mean values, each computed from the last 500 ns of one independent simulation.

#### Data Analysis

NUMPY, PANDAS were used for data analysis. MATPLOTLIB was used to generate plots. VMD^59,60^ was used to render figures for molecular visualizations.

#### Conformational landscape analysis

To quantify the overlap in conformational sampling, we combined all trajectories from our molecular dynamics (MD) simulations, including both the WT channel and its mutants (started from the up or down state conformations and using two different force fields), into a meta-trajectory summing up approximately 550 µs. The main chain atoms of the channel were extracted, and the trajectories were rotationally and translationally fitted to the minimized structures of the WT channel in its up or down state conformations. This fitting was performed using the S4-S2 binding sites of the selectivity filter as a reference.

We calculated the covariance matrix of the meta-trajectory and derived its eigenvalues and eigenvectors using the COVAR module implemented in GROMACS. The conformational sampling from conventional MD simulations was then projected onto eigenvectors 1 and 3, providing a low-dimensional representation of the sampled conformational landscape. Representative conformations along eigenvectors 1 and 3 were obtained using the ANAEIG module implemented GROMACS.

The probability density function (ρ) was derived from a 2D histogram of the conformations along eigenvectors 1 and 3, normalized by the sample count and bin area such that the total sum area equals 1. The free energy surface −Ln(ρ) was estimated as the negative natural logarithm of ρ, scaled so that the minimum value of the density corresponds to 0 K_b_T.

Contour plots were generated similarly to the probability density function ρ but were calculated separately for each mutant. A Gaussian kernel smoothing filter with a standard deviation of σ=7 was applied to the histograms. Each contour represents from 10% to 90% coverage of the total probability density.

The overlap coefficient was calculated by comparing the probability density functions of each mutant down state conformation with that of the WT up state conformation. The minimum values from these comparisons were integrated to define the overlap region and compute the total overlap coefficient. Confidence intervals (95 %) for the mean were estimated using bootstrapping.

#### Residue Interaction Networks

The sampled structures of TREK-2 WT and the mutants A317C, A318C, F215C, R237C, V319C, and W326C were used to analyse contacts between residues with the RING 3.0 web server^61^. The default parameters and distance thresholds were kept for the generation of the residue interaction networks (hydrogen bond: 3.5 Å, ionic/salt bridge: 4 Å, pi-cation: 5 Å, Van der Waals: 0.5 Å, pi-pi stacking: 6.5 Å, disulfide bond: 2.5 Å).

### Molecular biology and *in vitro* cRNA transcription

The coding sequences for the *rattus norvegicus* TREK-1b channel (*kcnk2* Isoform 1, GenBank accession number NM_172042.2), the human TREK-2a channel (*KCNK10* Isoform a, GenBank accession number NM_021161.5), and the human TRAAK channel (*KCNK4* Isoform 1, GenBank accession number NM_033310.2) were cloned into the pBF or pFAW vector. Point mutations were introduced by site-directed mutagenesis using custom primers containing the desired mutation and the obtained clones were verified by Sanger sequencing. Plasmid DNA was linearised with *NheI* or *MluI* restriction endonucleases and cRNA was transcribed *in vitro* using the SP6 or T7 AmpliCap Max High Yield Message Maker Kit (Cellscript, Madison, USA). cRNA was stored in nuclease-free water at -80 °C until used for injection.

### Oocyte preparation, injection and patch Clamp recordings

Electrophysiological studies were performed using oocytes surgically removed from adult female *Xenopus laevis*. Ovarian lobes were removed from tricaine anesthetised frogs and treated with 2 µg ml^-1^ type II collagenase in OR2 solution (in mM: NaCl 82.5, KCl 2, MgCl_2_ 1, HEPES 5; pH 7.4) for 60 min. After washing in ND96 solution (in mM: NaCl 96, KCl 2, CaCl_2_ 1.8, MgCl_2_ 1, HEPES 5; pH 7.5), Dumont stage V-VI oocytes were isolated, defolliculated and manually injected with ∼50 nl cRNA diluted to 10 - 100 ng µl^-1^. Oocytes were incubated in test solution (in mM: NaCl 54, KCl 30, CaCl_2_ 0.41, Ca(NO_3_)_2_ 0.33, MgSO_4_ 0.82, NaHCO_3_ 2.4, Tris 7.7; pH 7.4) at 17 °C for 12 - 72 h prior to electrophysiological experiments.

#### Inside-out patch clamp

Macroscopic currents were recorded from excised giant patches in inside-out voltage clamp configuration at room temperature (∼21 °C). Patch pipettes were made from thick-walled borosilicate glass (Science Products GmbH, Hofheim, Germany) and polished to yield resistances of 0.3 - 0.4 MΩ. Standard extracellular solution contained (in mM) 120 KCl, 10 HEPES, and 3.6 CaCl_2_ (pH 7.4 adjusted with KOH/HCl). The intracellular bath solution contained (in mM) 120 KCl, 10 HEPES, 2 EDTA, 1 pyrophosphate. The pH was adjusted to pH 5, pH 7.4 or pH 8 with KOH/HCl as indicated in the experiment description (HEPES was exchanged for MES in pH 5 solutions). Substances were applied to the patch by a multi-barrel application system. For NFx dose-response relationships in TREK-2 channels, the extracellular solution contained (in mM): 4 KCl, 116 NMDG, 10 HEPES, and 3.6 CaCl_2_.,. Data was acquired with EPC10 amplifiers controlled by PatchMaster software (HEKA electronics, Lamprecht, Germany), sampled at 10 kHz and filtered to a final bandwidth of 2.9 kHz with Bessel filters. Currents were recorded with a continuous pulse protocol at the voltage given in the experiment description or with a ramp protocol from -80 mV to +80 mV and plotted at the voltage as indicated in the experiment description.

#### Cell culture and whole-cell recordings

Whole-cell currents were recorded from transiently transfected HEK293 cells in voltage clamp configuration at room temperature (∼21 °C). HEK293 cells (RRID:CVCL_0045, Sigma Aldrich) were cultivated in Dulbecco’s Modified Eagles Medium (DMEM) supplemented with 10 % FCS and penicillin/streptomycin (100 U ml^-1^/100 µg ml^-1^) in 5 % CO_2_ at 37 °C. Cells were regularly tested for mycoplasm contamination by PCR (Venor GEM OneStep, Minerva Biolabs) and cell identity was visually controlled by comparison to the ATCC database. After transfection with Lipofectamine2000 (Invitrogen, Waltham, Massachusetts) according to manufacturer’s instructions, cells were incubated for 16 - 18h and seeded onto glass coverslips. Patch pipettes were fabricated from thin-walled borosilicate glass (Science Products GmbH, Hofheim, Germany), polished, and had resistances of 1.2 - 1.8 MΩ. Data was acquired with an EPC10 amplifier controlled by PatchMaster software (HEKA electronics, Lamprecht, Germany), sampled at 5 kHz and filtered to a final bandwidth of 2.9 kHz with Bessel filters. Currents shown were recorded with continuous voltage protocol at 0 mV in physiological K^+^ gradients (extracellular solution, in mM: 135 NaCl, 5 KCl, 2 MgCl_2_, 2 CaCl_2_, 10 Glucose, 10 HEPES, pH 7.4; intracellular solution, in mM: 140 KCl, 2 mM MgCl_2_, 1 mM CaCl_2_, 2.5 EGTA, 10 HEPES, pH 7.3). Series resistance was compensated to at least 75 %. LPC and NFx were applied to the cells by a multi-barrel application system.

#### Substances

Norfluoxetine hydrochloride (NFx) and 1-oleoyl 2-hydroxy-*sn*-glycero-3-phosphate (18:1 LPA) were purchased from Cayman Chemical via Biozol Diagnostics (Eching, Germany) and stored as 50 mM respectively 10 mM stock solution in DMSO at -20 °C. Tetrapentylammonium chloride (TPenA), Phosphatidylinositol-4,5-bisphosphate (PIP_2_), 1,2-dioleoy-*sn*-glycero-3-phosphate (18:1 PA), arachidonic acid (AA) and L-α-Lysophosphatidylcholine from egg yolk (LPC; 16:0/18:0/18:1 ∼ 69/27/3 %) were purchased from Sigma-Aldrich (Merck KGaA, Darmstadt, Germany). TPenA was stored as 100 mM stock solution in water at -20 °C, AA, PIP_2_, PA, and LPC as 10 mM stock solutions in DMSO at -20 - -70°C. Substances were diluted in bath solution to the final concentrations stated in the experiment description.

#### Data analysis

Electrophysiological recordings were analysed with PatchMaster software (HEKA electronics, Lamprecht, Germany) and fitted and visualized using Igor Pro (WaveMetrics Inc., Portland, USA). Currents at pH_i_ 8 were corrected for leaks by subtracting the remaining current in 1 mM TPenA. Relative *P*_o_ was calculated as

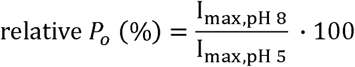

The maximal pH_i_ activated current was obtained by subtraction of background currents after full block by 1 mM TPenA. Relative *P*_o_ exceeded 100 % when a proton block masked the maximally activatable current at acidic pH_i_, and these values therefore represent apparent relative *P*_o_.

Dose-response relationships were fitted with a Hill equation

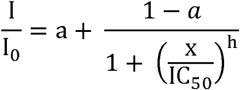

where I and I_0_ is the current with and without NFx/TPenA, *a* is the fraction of initial or residual current, x is the NFx/TPenA concentration, IC_50_ is the concentration at half maximal inhibition, and h is the Hill coefficient. All results are reported as mean ± SEM with n = number of experiments as stated in the figures or figure legends.

### Statistics

Statistical tests for electrophysiological data were carried out in GraphPad Prism version 8.4.3 for Windows (GraphPad Software, Boston, Massachusetts USA). Relative *P*_o_ results (Fig. 1d) were log transformed and tested for normal distribution. The Shapiro-Wilk test (α = 0.05) showed that the relative *P*_o_ measurements for 15 mutants did not have a log-normal distribution. A Brown-Forsythe and Welsh one-way ANOVA was performed to compare the effect of different mutations on relative *P*_o_. Dunnett’s T3 test for multiple comparisons was used as post-hoc test (α = 0.01). For mutants that did not have a log-normal distribution, a Kruskal-Wallis test for relative *P*_o_ (H = 116.4, p<0.0001, α=0.05) with a Dunn’s multiple comparisons post-hoc test (α = 0.05) was performed. The conclusions were the same as from the Brown-Forsythe and Welsh ANOVA for all mutants. Other tests used are described in the figure or table legends for the respective experiments.

All molecular dynamics simulations were performed using independent replicas of TREK- in either the up or down state, considering both WT and mutant systems and both force fields. The reported results correspond to averages over independent replicas. For the overlap parameter, confidence intervals were estimated by bootstrapping the mean.

### Ethics Statement

Animal husbandry and operation procedures of *Xenopus laevis* frogs used in this study comply to German animal welfare regulations and were certified by the authorities. Experiments using *Xenopus* frogs were approved by the local ethics commission of the Ministerium für Landwirtschaft, ländliche Räume, Europa und Verbraucherschutz (IX 555 – 106759/2023 (33-6/23 V)).

## Data availability

Crystallographic data of the protein structures used in simulations and for data visualization are available from the Protein Data Bank (PDB ID 4BW5 and 4XDJ for the human TREK-2 channel in up and down state, respectively, and PDB ID 4XDK for the NFx complex)^7^.

All analysed experimental and computational data (mean values given in the main text and used for figures) is enclosed as Supplementary Information (Tables S1 – S12).

The source data and computational results have been deposited in the GitHub https://github.com/EdwardMendez95/TREK2_updown.

## Code availability

The code used to estimate the sampled PCA space, quantify the degree of overlap, and generate the corresponding plots is available at https://github.com/EdwardMendez95/TREK2_updown. The dissipated work values used in the free-energy calculations are also provided in the same repository.

## Acknowledgements

We thank Prof. Stephen Tucker (University of Oxford, UK) for providing some of the TREK-1 mutants and we thank Petra Breiden, Michaela Unmack and Henning Janssen for excellent technical assistance.

## Abbreviations used

AA: Arachidonic Acid
2-APB: 2-aminoethyl-diphenylborinate
FEC: Free energy calculation
GOF: Gain-of-function
IC_50_: Concentration at half-maximal inhibition
K_2P_ channel: Two-pore domain K^+^ channel
LOF: Loss-of-function
LPA: Lysophosphatidic acid
LPC: Lysophosphatidylcholine
M2/M3/M4: Transmembrane helix 2/3/4
MD: Molecular Dynamics
NFx: Norfluoxetine
PA: Phosphatidic acid
PDF: Probability density function
PIP_2_: Phosphatidylinositol-4,5-bisphosphate
pH_i_: Intracellular pH
rel. *P*_o_: Relative open probability
SF: Selectivity filter
TPenA: Tetrapentylammonium
TREK-1/-2: TWIK-related potassium channel 1/ 2
WT: Wildtype

