## Supplementary Information for "Down- to up-state transition is the default pathway in TREK K_2P_ channel activation and does not involve a lipid occluded pore"

to

Supplementary Tables S1-S12

Supplementary Figures S1-S4

**Supplementary Table S1**: TREK-1 residues that exhibited relative open probabilities different from the wildtype and four wildtype-like control residues used for further analysis. A non-directional Brown-Forsythe and Welsh one-way ANOVA ($F = 36.45$, $p = <0.0001$; $W = 55.89$, $p = <0.0001$, $\alpha = 0.01$) with Dunnett's T3 test for multiple comparisons as post-hoc test ($\alpha = 0.01$) was performed to compare assay results from mutants and wildtype. Data represent mean ± SEM. TREK-2 residue codes are given for orientation.

| TM | TREK-1 | TREK-2 | Basal current $pH_i$ 8 (nA) | Rel. open probability (%) | P value (adjusted) | Phenotype |
|---|---|---|---|---|---|---|
| | WT | WT | 0.4 ± 0.1 | 4.8 ± 0.7 | - | |
| | L188C | L203 | 16.9 ± 3.0 | 87.1 ± 7.0 | <0.0001 | GOF |
| | L189C | L204 | 18.9 ± 3.3 | 61.6 ± 4.0 | <0.0001 | GOF |
| | A190C | A205 | 0.12 ± 0.02 | 1.6 ± 0.3 | 0.0038 | LOF |
| | G193C | G208 | 5.8 ± 1.3 | 33.8 ± 5.3 | 0.0004 | GOF |
| **M2** | G197C | G212 | 21.0 ± 1.9 | 100.6 ± 2.5 | <0.0001 | GOF |
| | T198C | T213 | 28.6 ± 6.0 | 55.2 ± 9.3 | 0.0038 | GOF |
| | I199C | I214 | 29.3 ± 7.1 | 87.8 ± 11.7 | 0.0018 | GOF |
| | F200C | F215 | 18.9 ± 4.6 | 110.3 ± 3.7 | <0.0001 | GOF |
| | E208C | E223 | 37.9 ± 5.6 | 76.3 ± 7.4 | 0.0003 | GOF |
| **M3** | R222C | R237 | 31.0 ± 3.3 | 93.1 ± 4.9 | <0.0001 | GOF |
| | I223C | V238 | 18.68 ± 3.35 | 33.0 ± 4.6 | 0.0004 | GOF |
| | W290C | W306 | 23.6 ± 3.8 | 96.7 ± 4.1 | <0.0001 | GOF |
| | L294C | L310 | 1.48 ± 0.83 | 16.6 ± 7.9 | 0.9917 | WT-like |
| | L297C | L313 | 5.8 ± 1.8 | 50.5 ± 3.1 | <0.0001 | GOF |
| | A301C | A317 | 0.0 ± 0.0 | 2.5 ± 0.4 | 0.0534 | WT-like |
| **M4** | A302C | A318 | 14.4 ± 2.8 | 121.9 ± 4.4 | <0.0001 | GOF |
| | V303C | V319 | 0.6 ± 0.2 | 6.0 ± 2.1 | >0.9999 | WT-like |
| | S305C | S321 | 0.6 ± 0.2 | 19.9 ± 4.6 | 0.2012 | WT-like |
| | M306C | M322 | 8.4 ± 2.1 | 46.2 ± 5.6 | <0.0001 | GOF |
| | W310C | W326 | 11.2 ± 2.4 | 48.2 ± 5.5 | <0.0001 | GOF |
| **pCt** | E321C | E337 | 10.1 ± 2.4 | 104.3 ± 7.4 | 0.0011 | GOF |

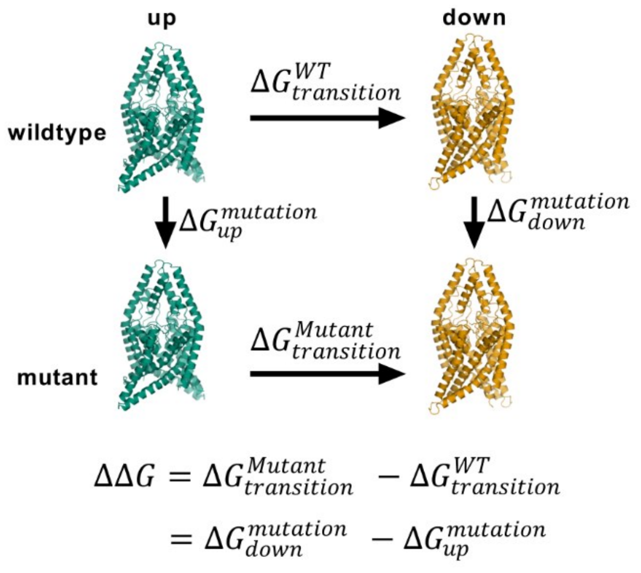


**Supplementary Figure S1**: Illustration of the thermodynamic cycle used in free energy calculations to determine ΔΔG values.

**Supplementary Table S2:** Free energy calculation (FEC) results and state-dependent Norfluoxetine (NFx) binding assay results for TREK residues that exhibited relative open probabilities different from the wildtype and for four wildtype-like control residues. A $\Delta\Delta G_{consensus}$ value of ≥ 4.2 kJ mol$^{-1}$ indicates destabilization of the down state, and 3x the NFx $IC_{50}$ of wildtype TREK-2 was used as threshold to assign the up conformation to a mutant. Data represent mean ± SEM. TREK-1 residue codes are given for orientation. n.d., not determined

| TM | TREK-1 | TREK-2 | $\Delta\Delta G_{consensus}$ (kJ mol$^{-1}$) | Conclusion FEC | NFx $IC_{50}$ (µM) | Conclusion Nfx |
|---|---|---|---|---|---|---|
| | WT | WT | 0.4 ± 0.1 | - | 3.65 ± 0.33 | down |
| M2 | L188 | L203C | -2.12 ± 0.48 | no conform. change | 2.98 ± 1.26 | down |
| | L189 | L204C | -3.76 ± 0.2 | no conform. change | 5.41 ± 0.78 | down |
| | G193 | G208C | -17.36 ± 1.05 | away from up | 8.62 ± 0.77 | down |
| | G197 | G212C | 9.97 ± 0.45 | away from down | 7.84 ± 1.87 | down |
| | T198 | T213C | -3.04 ± 0.43 | no conform. change | 3.09 ± 0.56 | down |
| | I199 | I214C | -1.98 ± 1.28 | no conform. change | 3.09 ± 0.58 | down |
| | F200 | F215C | 31.38 ± 1.2 | away from down | 33.08 ± 6.16 | up |
| | E208 | E223C | n.d | - | 12.39 ± 0.79 | down |
| M3 | R222 | R237C | n.d | - | 46.22 ± 11.80 | up |
| | I223 | V238C | n.d | - | n.d. | - |
| M4 | W290 | W306C | -6.04 ± 3.07 | away from up | 7.01 ± 1.03 | down |
| | L294 | L310C | -1.3 ± 1.18 | no conform. change | n.d. | - |
| | L297 | L313C | 3.78 ± 0.39 | no conform. change | 14.59 ± 1.52 | down |
| | A301 | A317C | -0.52 ± 0.66 | no conform. change | 3.15 ± 0.54 | down |
| | A302 | A318C | 23.84 ± 1.29 | away from down | 33.07 ± 2.97 | up |
| | V303 | V319C | 0.65 ± 0.58 | no conform. change | 7.23 ± 0.59 | down |
| | S305 | S321C | -2.68 ± 0.32 | no conform. change | n.d. | - |
| | M306 | M322C | 21.54 ± 0.57 | away from down | 4.08 ± 0.51 | down |
| | W310 | W326C | 9.29 ± 2.04 | away from down | 38.13 ± 6.05 | up |
| | E321 | E337C | n.d. | no conform. change | 30.17 ± 3.65 | up |

**Supplementary Table S3**: Mean overlap between the conformational spaces of down- and up-state derived from principal component analysis of the main chain in conventional MD simulations (mutants where the NFx assay indicated the up-state in bold).

| | Residue | Mean | Lower CI | Upper CI |
|---|---|---|---|---|
| Up-state inducing mutants | **F215C** | 0.0427 | 0.0025 | 0.0025 |
| | **R237C** | 0.2044 | 0.0039 | 0.0041 |
| | **A318C** | 0.1856 | 0.0044 | 0.0044 |
| | **W326C** | 0.1014 | 0.0029 | 0.0031 |
| Conflicting mutants | G212C | 0.0204 | 0.0018 | 0.0018 |
| | **L313C** | 0.1167 | 0.0040 | 0.0042 |
| | M322C | 0.021 | 0.0020 | 0.0020 |
| WT/ controls | WT | 0.0455 | 0.0021 | 0.0021 |
| | A317C | 0.0294 | 0.0023 | 0.0022 |
| | V319C | 0.0229 | 0.0020 | 0.0022 |

**Supplementary Table S4**: Individual residue interactions for up state mutants and control residues compared to the WT down state (in brackets)

| | up state inducing mutants | | | | | WT-like controls | |
|---|---|---|---|---|---|---|---|
| | **F215C** | **R237C** | **L313C** | **A318C** | **W326C** | **A317C** | **V319C** |
| H-bond | 5 (5) | 3 (4) | 4 (4) | 4 (4) | 3 (3) | 4 (4) | 4 (4) |
| π-cation | - | 0 (2) | - | - | 0 (2) | - | - |
| π-π-stack | 0 (4) | - | - | - | - | - | - |
| VDW | 7 (12) | 6 (8) | 2 (6) | 12 (5) | 5 (8) | 3 (2) | 3 (3) |

**Supplementary Table S5**: Protein-wide residue interactions for up state mutants and control residues and the WT (intra-chain / inter-chain)

| | up state inducing mutants | | | | | controls / WT | | | |
|---|---|---|---|---|---|---|---|---|---|
| | F215C | R237C | L313C | A318C | W326C | A317C | V319C | WT down | WT up |
| H-bond | 387/21 | 388/21 | 397/18 | 387/21 | 387/21 | 387/21 | 387/21 | 387/21 | 390/13 |
| π-cation | 3/2 | 1/2 | 0/0 | 3/2 | 1/2 | 3/2 | 3/2 | 3/2 | 1/1 |
| π-π-stack | 31/13 | 35/13 | 31/15 | 35/13 | 35/13 | 35/13 | 35/13 | 35/13 | 38/15 |
| ionic | 2/2 | 2/2 | 5/0 | 2/2 | 2/2 | 2/2 | 2/2 | 2/2 | 2/0 |
| VDW | 411/110 | 411/110 | 359/86 | 416/110 | 406/110 | 411/110 | 410/110 | 409/110 | 414/107 |

**Supplementary Table S6:** Free energy calculation (FEC) results and state-dependent Norfluoxetine (NFx) binding assay results for different substitutions at residue A318 in TREK-2. A $\Delta\Delta G_{consensus}$ value of ≥ 4.2 kJ $mol^{-1}$ indicates destabilization of the down state, and 3x the NFx $IC_{50}$ of wildtype TREK-2 was used as threshold to assign the up conformation to a mutant. Data represent mean ± SEM. TREK-1 residue codes are given for orientation.

| TREK-1 | TREK-2 | $\Delta\Delta G_{consensus}$ (kJ $mol^{-1}$) | Conclusion FEC | Nfx $IC_{50}$ (µM) | Conclusion Nfx |
|---|---|---|---|---|---|
| WT | WT | 0.4 ± 0.1 | - | 2.7 ± 0.4 | down |
| A302G | A318G | 7.51 ± 0.22 | away from down | 0.24 ± 0.07 | down + |
| A302C | A318C | 23.84 ± 1.29 | away from down | 33.07 ± 2.97 | up |
| A302V | A318V | 52.39 ± 0.62 | away from down | - | - |
| A302N | A318N | 61.19 ± 1.63 | away from down | - | - |
| A302F | A318F | 83.92 ± 2.4 | away from down | 155.73 ± 6.99 | up |
| A302Y | A318Y | 101.89 ± 3.02 | away from down | - | - |

**Supplementary Table S7**: TREK-1 residues that exhibited relative open probabilities different from the wildtype and four wildtype-like control residues used for further analysis. A non-directional Brown-Forsythe and Welsh one-way ANOVA ($F = 116.8$, p = <0.0001; $W = 311.5$, p = <0.0001, $\alpha = 0.05$) with Dunnett's T3 test for multiple comparisons as post-hoc test ($\alpha = 0.05$) was performed to compare assay results from mutants and wildtype. Data represent mean ± SEM. TREK-2 residue codes are given for orientation.

| TREK-1 | TREK-2 | Basal current $pH_i$ 8 (nA) | Rel. open probability (%) | P-value (adjusted) | Phenotype |
|---|---|---|---|---|---|
| WT | WT | 0.4 ± 0.1 | 4.8 ± 0.7 | - | |
| A302G | A318G | 0.04 ± 0.02 | 1.0 ± 0.6 | 0,0009 | LOF |
| A302C | A318C | 14.37 ± 2.80 | 121.9 ± 4.4 | <0,0001 | GOF |
| A302V | A318V | 2.02 ± 0.84 | 149.7 ± 7.0 | <0,0001 | GOF |
| A302N | A318N | 19.43 ± 3.24 | 133.2 ± 5.5 | <0,0001 | GOF |
| A302F | A318F | 7.71 ± 1.52 | 124.0 ± 6.9 | <0,0001 | GOF |
| A302W | A318W | 8.47 ± 4.66 | 71.6 ± 12.1 | 0,0011 | GOF |

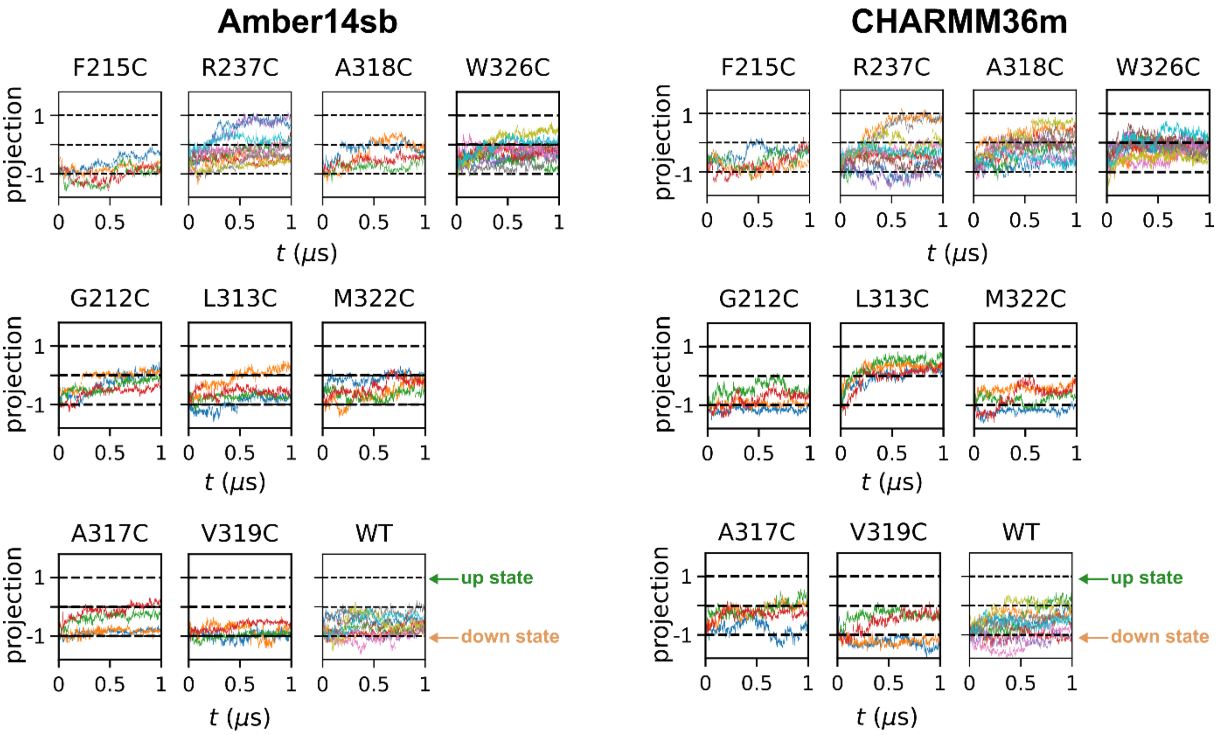


**Supplementary Figure S2**: Projection of the MD simulation trajectories onto the difference vector of the two crystallographic TREK-2 states for the WT, up state inducing cysteine mutants, and controls with both force fields used. A value of -1 represents the down conformation, and 1 represents the up conformation. A fluctuation about 0 in some simulations represents that either both subunits underwent a transition, or one subunit was in the up state while another subunit was in the down state. Simulations were started from the down conformation.

**Supplementary Table S8:** Mean distances d between M2 (Cα of G216) and M4 (Cα of W326) helices of TREK-2 wildtype and different substitutions at residue A318 derived from MD simulations.

| d down state | n | Mean (Å) | SEM (Å) | $\Delta$ distance to WT |
|---|---|---|---|---|
| WT | 40 | 11.76 | 0.40 | - |
| A318G | 16 | 10,01 | 0,53 | -1,75 |
| A318C | 28 | 14.97 | 0,56 | 3.21 |
| A318V | 16 | 16,11 | 0,64 | 4.35 |
| A318N | 16 | 17,06 | 0,83 | 5.30 |
| A318F | 16 | 15,62 | 0,62 | 3.87 |
| A318Y | 16 | 16,38 | 0,47 | 4.62 |
| **d up state** | | | | |
| WT | 40 | 20.49 | 0.18 | - |
| A318G | 16 | 20.79 | 0.23 | 0.30 |
| A318C | 28 | 19.91 | 0.36 | -0,58 |
| A318V | 16 | 21.17 | 0.29 | 0.68 |
| A318N | 16 | 20.91 | 0.33 | 0.42 |
| A318F | 16 | 19.95 | 0.46 | -0.55 |
| A318Y | 16 | 20.15 | 0.46 | -0.34 |

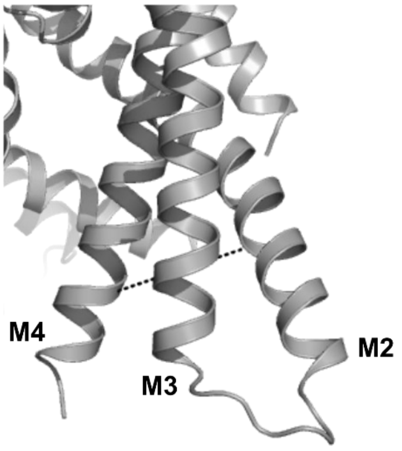

**Supplementary Table S9:** Mean $d_f$ (fenestration) distances between M2 and M4 helices of TREK-2 wildtype and different substitutions at residue A318 derived from MD simulations.

| | | $d_f$ | | |
|---|---|---|---|---|
| | n | Mean (Å) | SEM (Å) | Δ distance to WT |
| WT | 40 | 8.14 | 0.26 | - |
| A318G | 16 | 8.81 | 0.39 | 0.08 |
| A318F | 16 | 7.22 | 0.32 | -1.50 |

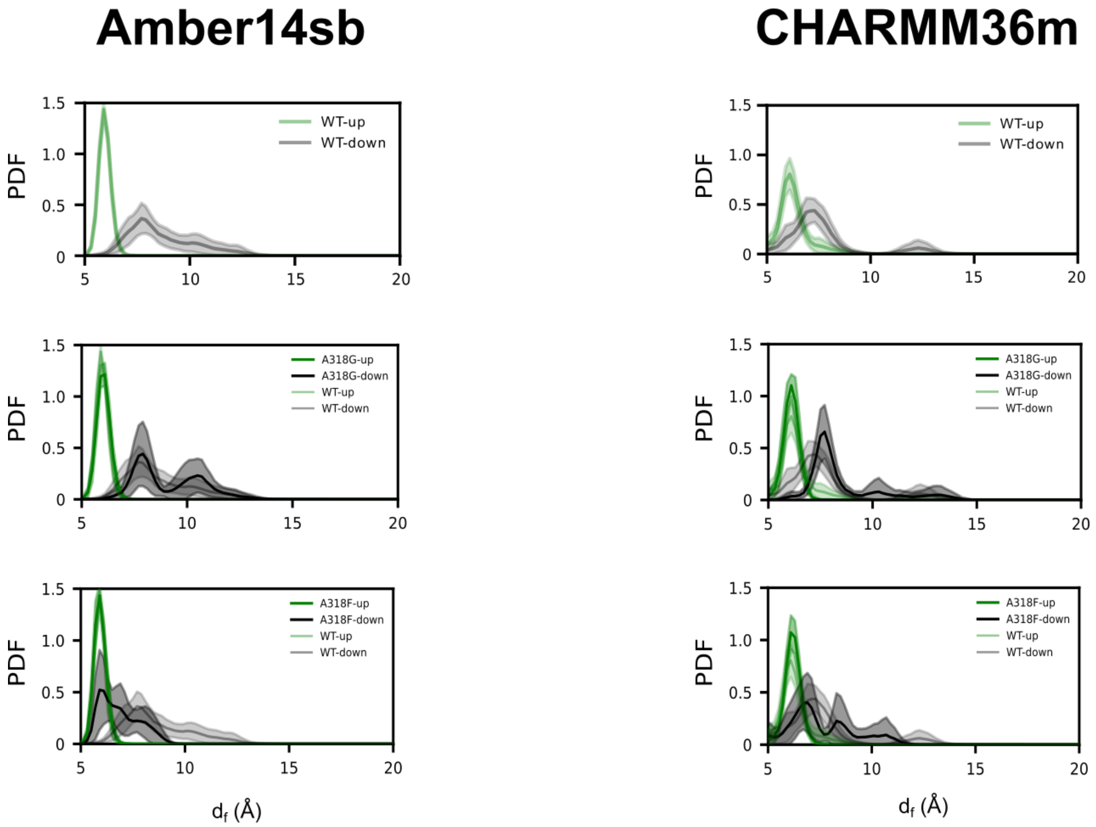


**Supplementary Figure S3**: Probability density functions of fenestration distances $d_f$ for WT A318G, and A318F substitutions with Amber14sb and CHARMM36m force fields used in MD simulations. Data are presented with 95 % confidence bands.

**Supplementary Table S10:** NFx $IC_{50}$ derived from measurements of dose-response relationships in TREK-2 channels activated by different stimuli. Data represent mean ± SEM.

| TREK-2 | NFx $IC_{50}$ (µM) | Conclusion |
|---|---|---|
| basal | 2.73 ± 0.36 | - |
| pHi 5 | 13.9 ± 1.13 | up state |
| LPA | 45.23 ± 8.10 | up state |
| PA | 54.24 ± 10.84 | up state |
| PIP2 | 38.35 ± 12.60 | up state |
| LPC | 12.2%* | up state |

**as mean difference with/without Nfx measured in whole cells (HEK293)*

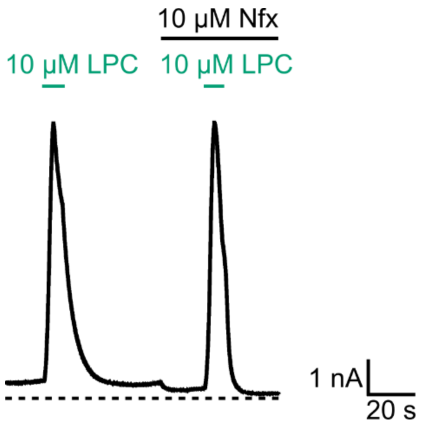


**Supplementary Figure S4:** Exemplary recording of TREK-1 channel activation by extracellular application of 10 µM LPC in the absence and in the presence of 10 µM NFx. Data were recorded in transiently transfected HEK293 cells at 0 mV with a continuous voltage protocol in physiological potassium gradients at pH 7.4.

**Supplementary Table S11:** TPenA $IC_{50}$ derived from measurements of dose-response relationships in TREK-2 wildtype, an up-state activatory mutant (A318C), and a down-state activatory mutant (I214C). A non-directional Brown-Forsythe and Welsh one-way ANOVA (F = 0.038, p = 0.9626; W = 0.035, p = 0.9653, α = 0.05) with Dunnett's T3 test for multiple comparisons as post-hoc test (α = 0.05) was performed to compare assay results from wildtype and mutant channels. Data represent mean ± SEM.

| TREK-2 | TPenA $IC_{50}$ (µM) | Adjusted P value |
|---|---|---|
| WT | 19.99 ± 4.93 | - |
| A318C | 12.39 ± 3.24 | ns |
| I214C | 25.72 ± 9.11 | ns |

**Supplementary Table S12:** Mean Nfx inhibition (%) of TREK-2 down state activating mutants before and after application of 10 µM $PIP_2$. A paired two-tailed t-test with the Holm-Sidak correction for multiple comparison was performed to compare inhibition in the basal and $PIP_2$ activated state (α = 0.05). Data represent mean ± SEM.

| TREK-2 | NFx inhibition (%) | | Adjusted P value |
|---|---|---|---|
| | basal | + $PIP_2$ | |
| T213C | 85.37 ± 5.24 | 19.60 ± 6.62 | <0.001 |
| I214C | 75.32 ± 5.66 | 26.11 ± 5.87 | <0.001 |
| W306C | 83.88 ± 6.95 | 34.69 ± 11.79 | 0.01 |

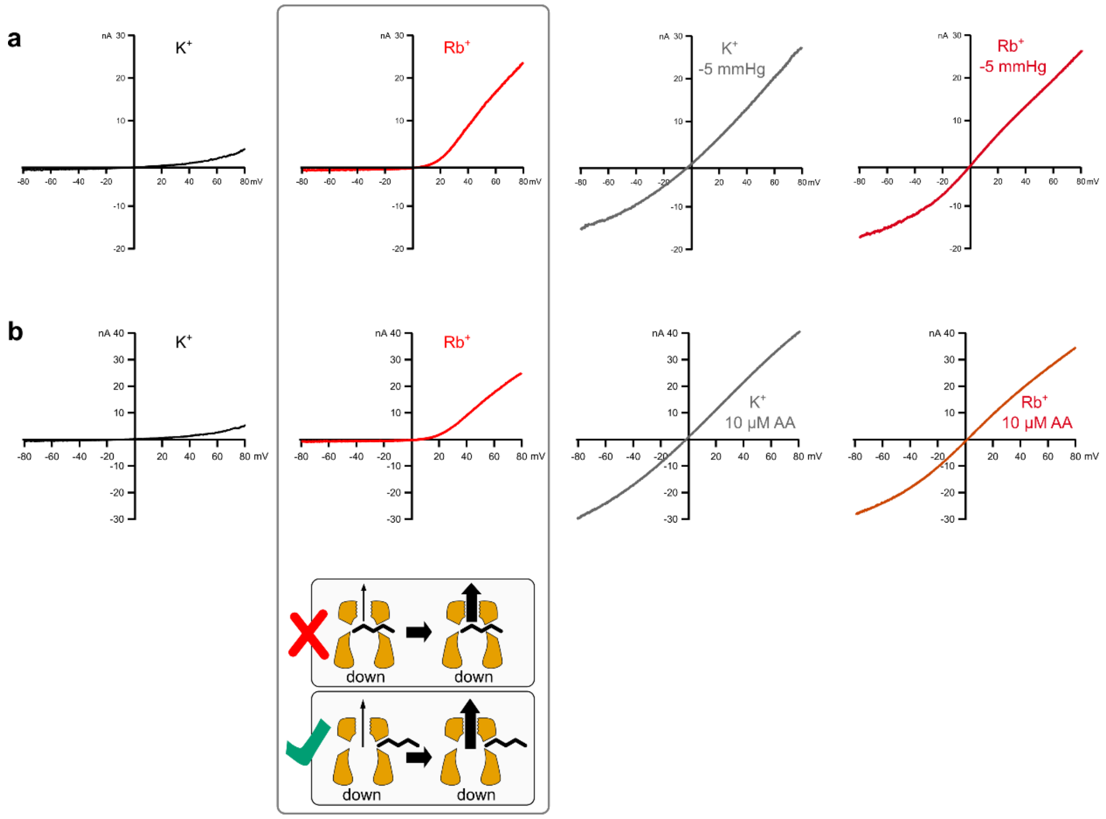


**Supplementary Figure S5**: Rb⁺ activation of TRAAK channels in the basal and activated state. **a** Exemplary ramp traces of TRAAK channels measured in K⁺, in Rb⁺, in K⁺ during stretch activation via negative pressure, and in Rb⁺ during stretch activation via negative pressure. **b** Exemplary ramp traces of TRAAK channels measured in K⁺, in Rb⁺, in K⁺ after activation by 10 µM AA, and in Rb⁺ activation by 10 µM AA. The comic depicts the concept that the observed Rb⁺ activation of the SF is not compatible with a lipid-blocked pore.
